# Mitotic chromosome compaction via active loop extrusion

**DOI:** 10.1101/021642

**Authors:** Anton Goloborodko, John F. Marko, Leonid A. Mirny

## Abstract

During cell division chromosomes are compacted in length by more than a hundred-fold. A wide range of experiments demonstrated that in their compacted state, mammalian chromosomes form arrays of closely stacked consecutive ∼100Kb loops. The mechanism underlying the active process of chromosome compaction into a stack of loops is unknown. Here we test the hypothesis that chromosomes are compacted by enzymatic machines that actively extrude chromatin loops. When such loop-extruding factors (LEF) bind to chromosomes, they progressively bridge sites that are further away along the chromosome, thus extruding a loop. We demonstrate that collective action of LEFs leads to formation of a dynamic array of consecutive loops. Simulations and an analytically solved model identify two distinct steady states: a sparse state where loops are highly dynamic but provide little compaction, and a dense state with more stable loops and dramatic chromosome compaction. We find that human chromosomes operate at the border of the dense steady state. Our analysis also shows how the macroscopic characteristics of the loop array are determined by the microscopic properties of LEFs and their abundance. When number of LEFs are used that match experimentally based estimates, the model can quantitatively reproduce the average loop length, the degree of compaction, and the general loop-array morphology of compact human chromosomes. Our study demonstrates that efficient chromosome compaction can be achieved solely by an active loop-extrusion process.

**Significance Statement:** During cell division chromosomes are compacted in length more than a hundred-fold and are folded in arrays of loops. The mechanism underlying this essential biological process is unknown. Here we test whether chromosome compaction can be performed by molecular machines that actively extrude chromatin loops. These machines bind to DNA and progressively bridge sites that are further and further away along the chromosome. We demonstrate that the collective action of loop-extruding machines can fold a chromosome into a dynamic array of loops. Simulated chromosome agrees with compact human chromosomes in its degree of compaction, loop size and the general loop-array morphology. Our study demonstrates that efficient chromosome compaction can be achieved solely by such active loop-extrusion process.

## Introduction

During cell division, initially decondensed interphase human chromosomes are compacted in length by more than a hundred-fold into the cylindrical, parallel-chromatid metaphase state. Several lines of evidence suggest that this compaction is achieved via formation of loops along chromosomes (1, 2). First, chromatin loops of 80 – 120 kb have long been observed via electron microscopy (2–4). These observations served as a basis for the “radial loop” models of the mitotic chromosome (4) and are consistent with optical imaging data (5). Second, theoretical studies showed that compaction into an array of closely stacked loops could explain the observed shape, the mechanical properties and the degree of compaction of mitotic chromosomes (6–8). More recently, the general picture of mitotic chromosomes as a series of closely packed chromatin loops was supported by Hi-C experiments, which measure the frequency of physical contacts within chromosomes (9). The same study independently confirmed the ∼ 100 kb length of the chromatin loops.

The mechanism underlying compaction of chromosomes into a stack of loops is unknown. Several lines of evidence suggest that this compaction cannot be achieved by simple mechanisms of chromatin condensation, e.g., “poor solvent” conditions, or non-specific chromatin “cross-linker” proteins. First, the loops are formed overwhelmingly within individual chromatids. Different chromosomes and sister chromatids are not extensively cross-linked to each other as would tend to happen during nonspecific condensation, but instead become individualized during the compaction process. Second, loops are arranged in essentially genomic order and are non-overlapping (9), without the strong overlap of loops that would be expected from nonspecific crosslinking. Finally, metaphase chromosomes compact into elongated structures with a linear arrangement of loops along the main axis. A cross-linking agent would generate surface tension and shrink chromosomes into spherical globules with a random spatial arrangement within a globule (6, 10, 11). In fact, the term “condensation”, which generally refers to the effects of chemical interactions driving phase separation and surface tension, is inappropriate for description of mitotic chromosome compaction where neither effect occurs. Chromatin is clearly being actively compacted during mitosis.

An alternative hypothesis is that chromosomes are condensed by enzymatic machines that actively extrude chromatin loops (12, 13). When these enzymes bind to chromosomes, they first link two adjacent sites, but then move both contact points along the chromosome in opposite directions, so that they progressively bridge more distant sites (12). Loop-extruding functions have been observed for other enzymes acting on naked DNA (14–17). Condensin complexes, which play an central role in chromosome compaction (18) and which are present in all domains of life (19), are likely to be a key component of such “loop-extruding factors” (LEFs). A key question is whether LEFs alone are sufficient to drive formation of arrays of non-overlapping loops essential for linear compaction of chromatids, or if other factors are required, e.g., to define the loop bases.

In this paper, we model the collective action of loop-extruding factors (LEFs) that dynamically exchange between the nucleoplasm and chromatin fiber. We find that LEFs self-organize into a dynamic array of consecutive loops, which has two distinct steady states: a sparse state where loops are separated by gaps and provide moderate compaction, and a dense state where jammed LEFs drastically compact a long chromatin fiber. These states can be described by a simple analytical model of loop dynamics. We show how the macroscopic characteristics of the loop array are determined by the microscopic properties of the LEFs and their abundance, and we demonstrate that efficient chromosome compaction can be achieved solely by LEFs.

## Results

### Model for LEFs on a long chromatin segment

To understand the dynamics of loops formation and chromatin compaction by loop-extruding factors (LEFs) we carried out stochastic simulations of the process shown in Fig. 1 (13). We focus on the organization and dynamics of loop formation and dissolution without considering 3D organization of the chromatin fiber and assume that emerging topological conflicts can be resolved by topoisomerase II enzymes active during metaphase compaction.

**Figure 1.**
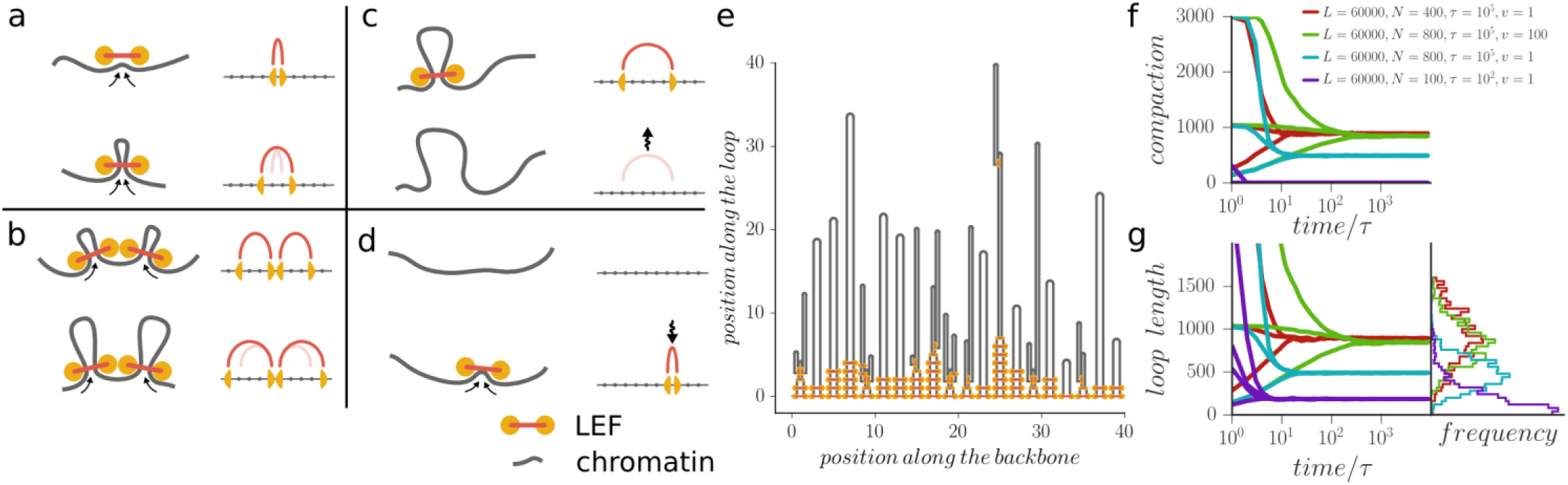
Simulations of chromosome compaction by loop extruding factors (LEFs). The action of LEFs can be simulated using a one-dimensional lattice model with four dynamic rules (a)-(d): (a) LEFs extrude loops by moving the two connected heads along the chromosome, (b) LEF heads block each other, (c) LEFs dissociate from chromatin, and (d) LEFs in the solution rebind to the chromosome. (e) Simulations show that LEFs can fold a chromosome into an array of consecutive loops. The diagram shows the loops formed by LEFs in a simulation with *L* = 2000, *N* = 200, *τ* = 450, ν = 1 after 45000 time steps. (f),(g) The system of LEFs on a long chromatin fiber converges to a steady state. The steady distribution of loop sizes and the degree of compaction depends on the control parameters, but is independent of initial state. Results are shown for different simulation parameters and starting conditions; data for each curve is averaged over 10 simulation replicas.

We consider a single piece of chromatin fiber of length *L*, occupied by *N* LEFs. We model a LEF very generally as having two “heads” connected by a linker. The LEF heads bind to nearby sites along the chromatin fiber and proceed to slide away from each other stochastically with an average velocity *v*, thus extruding a loop with rate *2v* (Fig. 1a). When the heads of two neighboring LEFs collide, they block each other and halt (Fig 1b).

For the LEFs to be able to organize robust loop domains, it is essential that they be able to relocate via unbinding and rebinding (13). We allow each LEF to dissociate at a rate1/*τ* which is independent of their state and location (Fig 1c). When a LEF dissociates, we suppose that it immediately rebinds to a random site elsewhere on the chromosome, where it begins to extrude a new loop (Fig. 1d). The model is fully determined by the four parameters (*L, N, ν, τ*) of which *N* and *τ* can be estimated from the experimental studies of condensins (20–22). We divide the chromosome into *L*=6.10^4^ sites, so that each site can be occupied by one LEF head. With each site roughly corresponding to a nucleosome with a DNA linker (∼ 200 bp or ∼10 nm, a fraction of a size of a condensin complex), our simulated chromosome corresponds to ∼ 12 Mb of chromatin fiber.

### LEFs can generate a tightly stacked loop array and strong chromosome compaction

In initial simulations we observed that the LEFs generated tightly stacked loops with a high degree of chromatin compaction, despite their constant dissociation (Fig. 1e-g, Supplemental Movie 1). To test that this was a steady state rather than a frozen (glassy) configuration we performed simulations ten times longer than the apparent time needed to reach the steady state, and used a broad range of initial conditions (SI). Simulations converged to states with degree of compaction and distribution of loop size which depended on the control parameters, but were independent of initial states (Fig 1f,g and SI Appendix), providing further support to the existence of a well-defined loop-stacked steady state.

### Two characteristic lengths control whether LEFs form dense or sparse chromatin loops

To understand how the microscopic characteristics of the LEFs control the compaction process, we performed simulations systematically exploring the control parameter space. This revealed that there are two distinct steady states of loop-extrusion dynamics in the model (Fig. 2): i) a sparse, poorly compacted state where loops are formed by single LEFs and separated by gaps (Fig. 2c); and ii) a dense state, where the chromosome is compacted into an array of consecutive loops each having multiple LEFs at its base (Fig. 2d).

**Figure 2.**
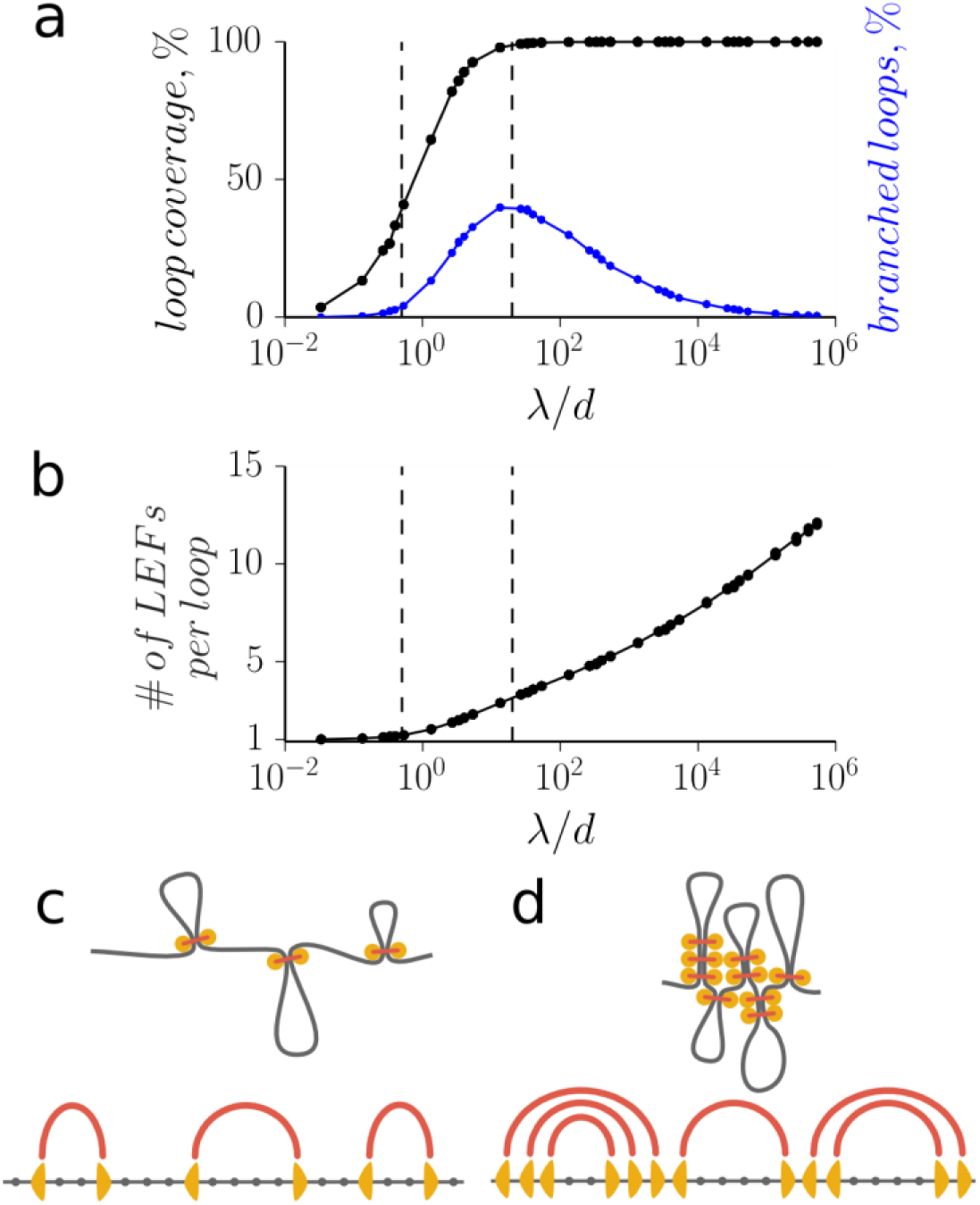
Simulations of LEFs reveal two distinct steady states. (a-b) The properties of loop arrays formed by LEFs, such as the portion of the chromosome extruded into loops, the portion of branched loops and the number of LEFs per loop, depend on the dimensionless ratio λ /*d*. This ratio defines the two steady states of the system: (c) the sparse state (λ /*d* ≪ 1) where the loops are supported by single LEFs and separated by big loop-free gaps and (d) the dense state (λ /*d* ≫ 1) where the whole chromosome is extruded into an array of consecutive loops supported by multiple LEFs. In both steady states, the loops are not branched (a). The vertical dotted lines at λ /*d =* 0.5 and 20 roughly show the transition region.

In the sparse state, LEFs do not efficiently condense chromosomes, since even a small fraction of fiber length remaining unextruded in the gaps between loops prevents efficient linear compaction (Figure S7). In the dense state, however, the whole chromosome is folded into a gapless array of loops, where the end of one loop adjacent to the beginning of the following one (Fig 2). Such organization was found to be essential to achieve agreement with Hi-C data for mitotic chromosomes (9). Below we show that realistic LEF abundance (one condensin per 10 – 30 kb) can give rise to loop sizes of ∼ 80 – 120 kb consistent with mitotic Hi-C and earlier direct measurements (3, 21–24) and inferred from Hi-C data (9). These findings suggest a dense state as an attractive model of chromosome compaction.

Two steady states arise from the interplay of two length scales characterizing the LEFs: i) *processivity λ* = *2ντ*, the average size of a loop extruded by an isolated LEF during its residency time on chromatin; and ii) the average linear *separation* between LEFs *d* =*L*/*N* (Fig 2a). When λ/*d «* 1, the system resides in the sparse state: LEFs work in isolation, a small fraction of the chromosome is extruded into loops and large gaps between them prevent efficient compaction. In the opposite dense case, when λ/*d » 1*, the whole chromosomal fiber is extruded into loops, leading to a high degree of compaction. When the loop coverage is plotted as a function of *λ*/*d*, rather than individual parameters, the curves collapse into a single transition curve indicative of the central role of *λ*/*d* in controlling the compaction (Figure S1)

### Loop organization and dynamics are distinct in the sparse and dense steady states

To understand the process of chromosome compaction by LEFs, we consider the dynamics of loop formation and disassembly. In the sparse state, LEFs rarely interact; each loop is extruded by a single LEF and it disappears once the LEF dissociates, leading to a highly dynamic state with a rapid (∼ *τ*) turnover of loops (Fig S1a, SI Appendix). Since LEFs extrude loops continuously and the distribution of LEFtheir residence times is exponential, the distribution of loop size is exponential too (Figure S2).

Two aspects of the loop organization control the dense state dynamics: i) loops have no gaps between each other; and (ii) individual loops are reinforced by multiple LEFs, i.e. several LEFs are stacked on top of each other at the base of a loop (13). Both phenomena result from the competition for chromosomal fiber among abundant LEFs. The gaps disappear because, in the dense state, LEFs have enough time to extrude all available fiber until colliding with adjacent LEFs. Abundant collisions lead to a non-exponential distribution of loop sizes (Figure S2, SI Appendix). Loop reinforcement is also caused by LEF collisions (Fig 3): every time a new LEF binds within an existing loop, it re-extrudes this loop until colliding with the LEF residing at the loop base. As a result, each loop is stabilized by multiple LEFs at its base. The absence of gaps and the reinforcement of loops preserve the structure of loops on the timescales *t » τ* (Fig S1b, SI Appendix): loops cannot grow because their LEFs are blocked by the neighbors, and they do not disband when individual LEFs dissociate, as remaining LEFs support them. Thus, the loops of a condensed chromosome are maintained despite continuously exchanging LEFs, like the planks in the ship of Theseus (25).

**Figure 3.**
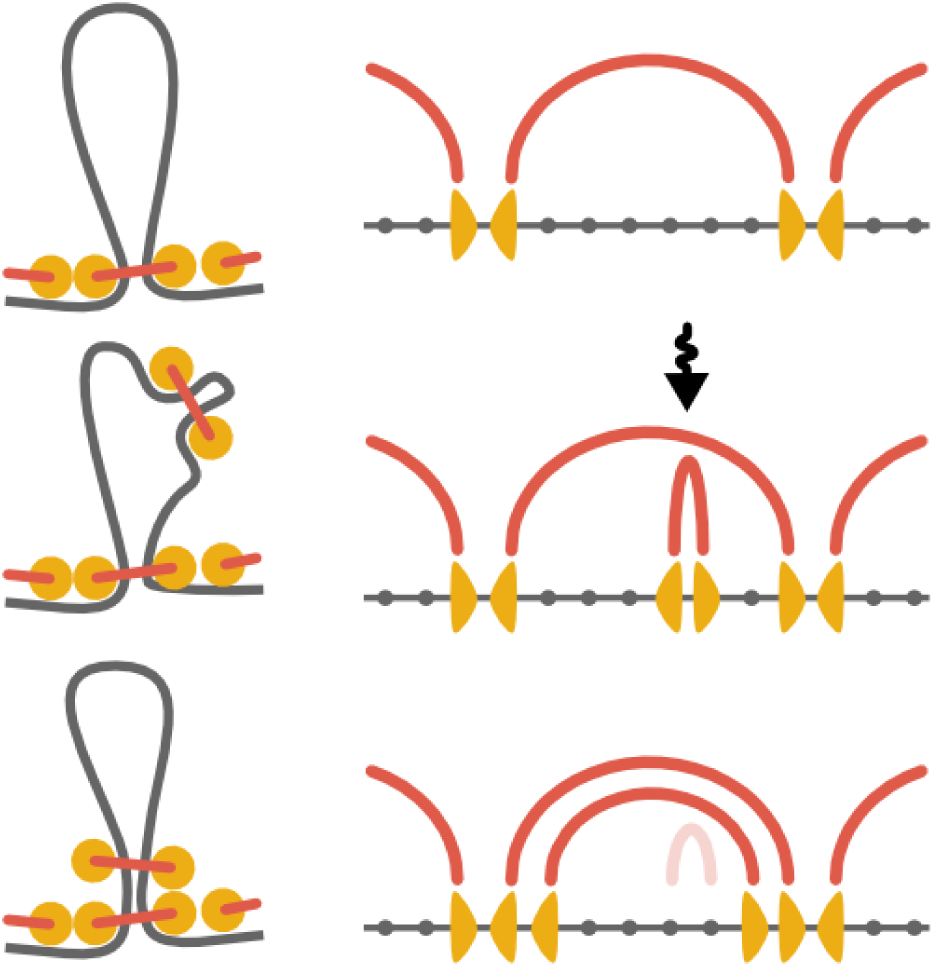
The mechanism of loop reinforcement in the dense state. Upon binding to an existing loop, a LEF reextrudes it and stacks on top of the LEFs already supporting the loop.

### Steady state loop dynamics is controlled by competition between loop death and division

To develop an analytical model of the system’s steady state we consider its dynamics. Loops in the dense state are not completely static: Two stochastic processes, loop “death” and loop “division”, change the structure of the loop array and drive self-organization of the steady state.

A loop “dies” when the number of LEFs at its base supporting it fluctuates to zero. When all LEFs dissociate, neighboring LEFs become unblocked and extrude the released fiber into their own loops (Fig 4a). We compute the rate at which a stack of *n* LEFs supporting a loop of size *ℓ* can stochastically fluctuate to zero. The stack can shrink due to LEFs dissociation (at rate *n*/*τ*) and can grow due to association of new LEFs to the loop (at rate 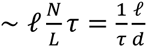). Fluctuations of the LEF stack size are equivalent to the stochastic immigration-death process, for which the rate of fluctuation to zero can be computed as 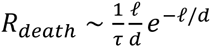 (SI Appendix) (26).

**Figure 4.**
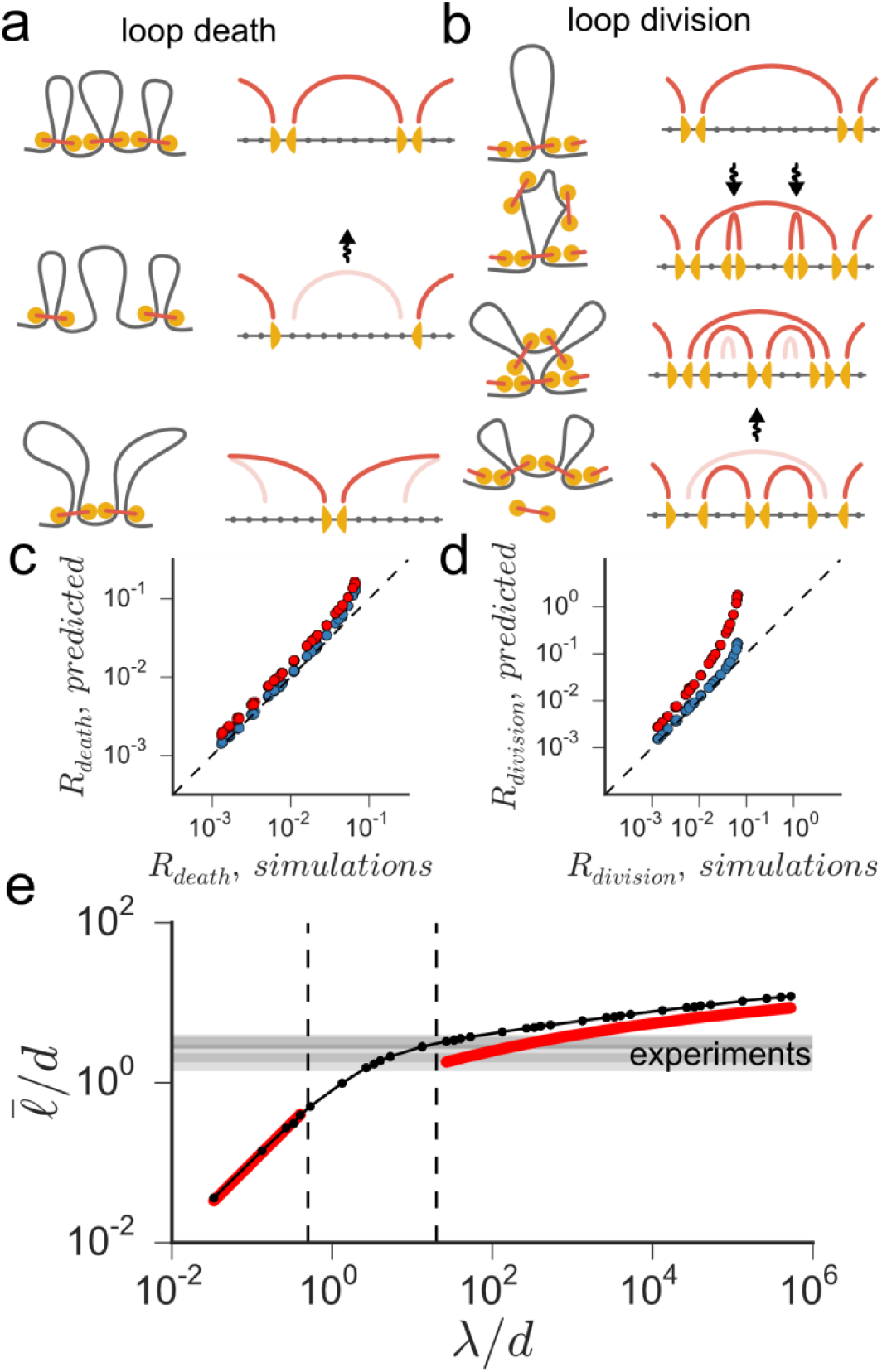
The model of loop death and division explains the origin of the dense steady state. (a) Loops occasionally disassemble when the number of reinforcing LEFs fluctuates to zero. The chromatin of the disassembled loop is immediately extruded into the adjacent loops. (b) A loop splits upon simultaneous landing of two reinforcing LEFs. The rates of loop death (c) and division (d) in the dense state can be estimated using simple analytical formulas (red dots) or more accurate computational models (blue dots). (f) In the dense state, the steady-state balance between loop death and division provides an approximate analytical expression for the average loop length (the red line). In the sparse state, the average loop length is predicted to be equal λ (the red line). Both predictions agree well with the simulations (the black line). The four horizontal overlapping gray bands show the available independent experimental estimates of 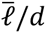 in mitotic human chromosomes: 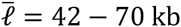, 54 – 112 kb (24), 80 –90 kb (23) and 80 – 120 kb (9) and *d* ≈ 30 kb (21).

A loop “divides” into two smaller loops when two LEFs land within a single loop almost simultaneously and extrude two smaller consecutive loops (Fig 4b). These newly created loops become subsequently reinforced by other LEFs that land into them. The original “parent” loop, on the contrary, is effectively cut off from the supply of reinforcing LEFs, and disintegrates on a timescale ∼ *τ*, with the two child loops taking its place. The rate-limiting process for loop division is the landing of two LEFs onto the same loop, giving an estimate for the rate of division: 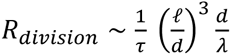 (SI Appendix). These scaling laws accurately predict the dynamics of loop birth and death (Fig. 4c, d)

In the steady state, the number of loops is approximately constant. By equating the rate of loop creation by division to the rate of loop death, the average loop size is obtained as:

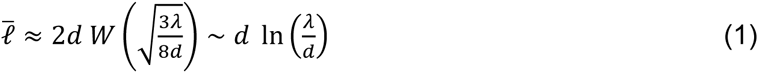

and the average number of LEFs per loop is:

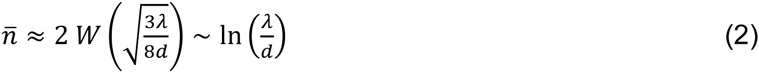

where *W*(*x*)is the Lambert W function.

Our analytical model agrees with simulations (Fig 4e) and explains how the number of LEFs and their microscopic properties affect the morphology of compacted chromosomes. First, Eq. (1) and (2) show that λ/*d* is the key control parameter of the system, which determines not only the state of the system (sparse vs dense), but also loop sizes and the degree of loop reinforcement in the dense state.

Using these scaling laws plus available experimental data we can estimate LEF processivity and dynamic state for human metaphase chromosomes. The average loop length has been estimated by miroscopy and via modeling of Hi-C data as *ℓ* = 80 – 120 kb (3, 9, 23, 24). The spacing between bound condensin molecules was measured as *d* = 30 kb (21). Using these values we obtain a range of *ℓ*/*d* ≈ 3 that is shown on Fig 4e and corresponds to *λ*/*d* ≈ 20. These values shows that human mitotic chromosomes operate at the lower bound of the dense state, have each loop reinforced by n ≈ 3 LEFs, and human LEF have processivity λ ≈ 600 kb.

Second, our analysis allows to compute the degree of chromosomal compaction by LEFs. Since the length of a compacted chromosome in the gapless dense state equals the sum of the widths of the loop bases, *a* (Fig 1E), the coefficient of chromosomal compaction is 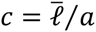. While addition of extra LEFs leads to better loop reinforcement, it also makes loops shorter (*l ∼ ln N*/*N*) and thus reduces the degree of chromosomal compaction (Fig S7, SI Appendix). For the loop base size close to the chromatin fiber diameter a = 10-20 nm we obtain the degree of compaction 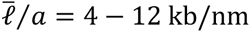 Interestingly, our estimate for the compaction achieved through folding of a chromosome into a gapless array of loops is in a good agreement with the experimentally measured degree of human chromosome compaction in mid-prophase (∼ 6 kb/nm) (27).

Third, our model predicts how the loop array morphology changes in response to biological perturbations. Specifically, factors that decrease the speed of loop extrusion *v* or reduce LEF residence time *τ* will decrease the processivity λ and thus decrease the average loop size, degree of loop reinforcement and the degree of chromatid compaction. The effects of LEF overexpression depend on the state of the system: for sparse loop arrays, it does not affect the average loop size and only increases the number of loops and, thus the degree of compaction. In the dense systems, LEF overexpression decreases the average loop size and degree of compaction, but increases the degree of loop reinforcement.

Finally, this analytical model shows how LEFs robustly self-organize chromosomes into a globally stable steady state. The rates of death and division *R*_*death*_ and *R*_*division*_ scale differently with the loop length: large loops are more likely to divide into smaller ones (*R*_*death*_ ∼ *ℓ*^3^), and smaller loops are more likely to die (*R*_*death*_ ∼ *ℓe*^−ℓ^) allowing neighboring loops to grow. This negative feedback drives the system to a steady state with a relatively narrow distribution of loop lengths. These results indicate that loop sizes and hence chromosome diameter and length will be sensitive to concentrations of LEFs while the overall morphology as a gapless array of consecutive loops will remain unchanged as long as the system remains in the dense state.

## Discussion

Our model of loop-extrusion provides a resolution of the puzzle of how roughly nanometer-sized enzyme complexes can drive the regular organization of a chromosome at scales well beyond a micron, as occurs in eukaryote cells during mitosis. A fundamental problem with almost any mechanism based on non-specific crosslinking of chromatin fibers is that chromosomes will end up crosslinked together: our model avoids this fate by having LEFs bind to chromatin at one location and then actively extrude loops without the possibility of forming inter-chromosome attachments. Through unbinding, rebinding, and re-extrusion, enzymatic machines of this type gradually build larger loops anchored by multiple LEFs, eventually reaching a steady state. A key feature of our model is that the compaction process proceeds by a combination of stochastic loop “death” and “division” events, which gradually but not strictly monotonically leads to a highly compacted chromosome.

Compaction driven by LEFs is distinct from the usual polymer “condensation” occurring under “poor solvent” conditions. Unlike proposed linear compaction, nonspecific adhesion of chromatin fibers to one another would generate surface tension, driving adhesion of chromosomes together into spherical masses of chromatin (11), increasing entanglement and working against chromosome segregation and individualization (8, 13). An important feature of the LEF-compacted state is that despite its robust structure, it is entirely dependent on DNA connectivity; intermittent cleavage of DNA alone can lead to dissolution of the entire chromosome, as has been observed experimentally (28, 29).

We emphasize that in the compacted steady state the loops have a well-defined size, and that inside the chromosome the LEFs establish internal tension, rather than the surface tension generated by nonspecific crosslinking. This internal tension is an essential contributor to the uniform folding and well-regulated cylindrical morphology of chromatids, and also generates repulsive forces between folded chromosomes essential to segregation of sister chromatids and individualization of different chromosomes (6).

Note that achieving a compacted steady state by this mechanism can be a slow process, i.e. it would require ∼10^2^ –10^3^ *τ*, while the turnover rate *τ* for condensin was measured to be at least a few minutes (20). However, we found that gradual or step-wise loading/activation of LEFs can lead to a significant speed-up of the process (SI Appendix). For the optimal activation rate, the compact steady state can be achieved in a fraction of time *τ* (see Fig S9, SI Appendix). A sign of this dynamics occurring *in vivo* would be its gradual upregulation/activation of condesin during early chromosome compaction.

On the other hand, if there are too few LEFs we have found that a distinct, disordered, poorly compacted chromosome steady state occurs. This outcome has been observed in experiments where condensins were interfered with, both in cells (30) and in *Xenopus* egg extracts (18, 30). Modulation of chromosome structure also has been observed to occur through development, for example in *Xenopus*, where mitotic chromosomes become gradually shorter and fatter with maturation (31); this gradual change in chromosome morphology could be due to changes in LEF amount or activity with development.

While the mechanisms of loop extrusion remain unknown, a relative simple molecular organization of a protein complex could produce loop-extruding activity. A LEF composed of two connected heads, each able to move along chromatin fiber processively, can achieve a loop extrusion activity. Moreover relative dynamics of the two heads (motors) does not have to be coordinated. In fact, of four possible relative orientation of heads’ directions (→→, ←← →←, ←→), two (→→, ←←) produce LEFs that slide along chromatin without loop extrusion, one (→←) makes LEFs with heads pushing against each other and thus stuck on chromatin, and the last one (←→) makes LEFs with heads moving away from each other and thus extruding loops. It remains to be seen how these functions are implemented in structures of SMC complexes (cohesins and condensins) that have enzymatic (ATP-hydrolyzing) domains.

We note that our model does not discern between condensin I and condensin II, and also that the considered compaction process is the prophase compaction driven by condensin II (30). Experiments aimed at disrupting LEFs would perhaps be best targeted at condensin II; however other proteins may be involved as condensin II by itself is not thought to have motor function. Intriguingly, the motor KIF4A has been shown to be involved with mitotic chromosome compaction (32); it is conceivable that condensins are somehow aided in a LEF function by a separate motor molecule such as KIF4A. Alternately, condensins may be able to cooperatively organize so as to generate contractile LEF behavior, for example by “directional polymerization” (13).

## Materials and methods

Simulations were performed using the Gillespie algorithm (13, 33). The Python code performing the simulations of loop extrusion and the data analysis and is available online at https://bitbucket.org/golobor/loop-extrusion-1d. See SI Appendix for details of simulations.

## Acknowledgements

Work at NU was supported by the NSF through Grants DMR-1206868 and MCB-1022117, and by the NIH through Grants GM105847 and CA193419. Work at MIT was supported by the NIH through Grants GM114190 R01HG003143. We are grateful to Maxim Imakaev, Geoffrey Fudenberg, Christopher McFarland, Nezar Abdennur, Rotem Gura, Mehran Kardar and the members of MIT Biophysics for productive discussions.

## Abbreviations

LEF – loop extuding factor

SMC – structural maintenance of chromosomes

## Supplementary Information Appendix

### Mitotic chromosome compaction via active loop extrusion

#### 1 Materials and methods

We study the action of loop extruding factors (LEFs) using the previously described model [1]. In this model, we model a chromosome as a one-dimensional lattice with *L* sites with *N* LEFs. Each LEF is represented as a pair of “heads”, each occupying an individual site on the chromosome. The positions of LEF heads are stochastically updated using the Gillespie algorithm with four rules:

1. The two heads of each LEF stochastically step away from each other with the average rate *v*.
2. The heads of different LEFs cannot step over each other and thus stop extrusion upon reaching another LEF. However, the two heads of the same LEF extrude loops independently and if one head of a LEF is blocked, another head continues extrusion.
3. LEFs stochastically unbind from the fiber with the rate of 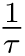, where *τ* is the average residence time.
4. Free LEFs immediately rebind to the chromatin fiber at a random uniformly chosen pair of adjacent sites.

In this study we modeled 12Mb of chromatin fiber, close to the size of the smallest human chromosomal arm 21p (12.7 Mb). Without loss of generality, we divided the fiber into a lattice of *L* = 60000 cells of 200 bp each, roughly the size of a nucleosome with a DNA linker. We simulated systems with *N* = 100, 400, 800, 1200 and 1600 LEFs, where 400-1200 LEFs corresponded to the experimental estimates of the abundance of condensin in mitotic human cells (1 per 10-30kb) [2, 3]. The speed of extrusion *v* varied in a broad range between 1 and 100 sites per time unit and the residence time *τ* varied between 10 and 10^5^ time units. At the beginning of each simulation, LEFs were distributed randomly along the chromosome with both heads in adjacent lattice sites. We simulated each system for 10^4^ *τ* units of time, with 10 simulations per each parameter set.

We found that the average loop length *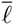* in the steady state did not depend on the initial positions of LEFs, with only 1 out of 75 tested parameter sets failing the Bonferroni-corrected one-way ANOVA comparison of 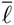 between ten randomly initiated replicas. Additionally, we found that the same final values of 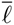 were achieved if the system was initiated with 20 or 60 loops of equal length, each supported by closely stacked LEFs.

#### 2 The two regimes of LEFs on a chromosome

We found that the system of LEFs has two distinct regimes: the sparse regime, where loops are formed by individual LEFs and are separated by gaps, and the dense regime, where loops are supported by multiple LEFs and cover the chromosome completely. The transition between the two regimes occurs when we increase the parameter 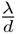, where *λ* = 2*vτ* is the LEF *processivity*, i.e. the average length of a loop extruded by an obstructed LEF over its residence time on chromatin and 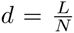 is the average linear *separation* between LEFs. Our simulations suggested that this ratio, and not each of the parameters alone, determines the average loop coverage, i.e. the portion of chromatin extruded into loops (Figure S1).

**Figure S1:**
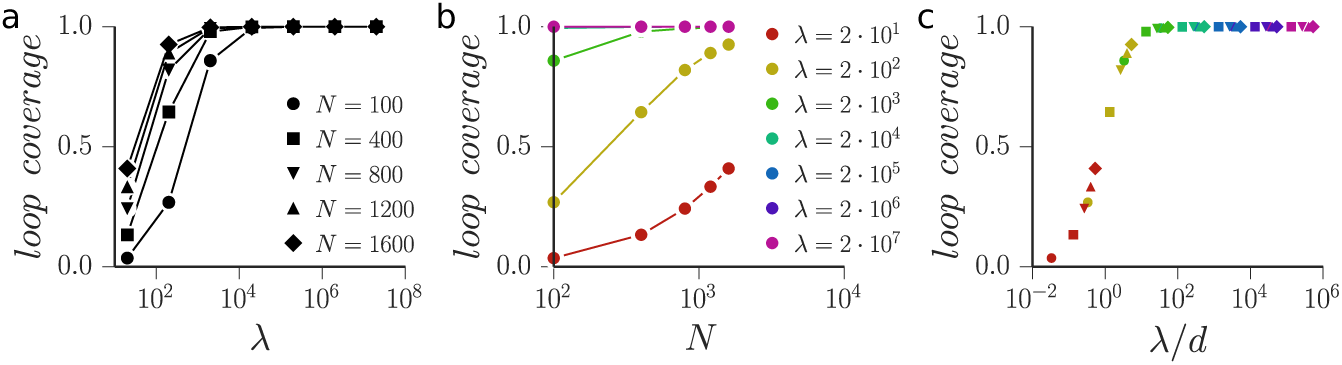
Loop coverage as (a) a function of the LEF processivity *λ* for different numbers of LEFs *N* and (b) as a function of *N* for several values of *λ*. (c) The curves collapse when plotted relative to the ratio *λ*/*d* = *λN* /*L*.

##### Loops have different dynamics in the sparse and dense regimes

We found that loops in the two regimes of LEFs display very different dynamics (Fig. S2). As a proxy for the timescale of loop stability we measured the autocorrelation time of the LEF footprint on chromatin. This measure allowed us to estimate the characteristic times of change in loop structures across multiple orders of magnitude. In the sparse regime, autocorrelation time is much shorter than *τ*, inversely proportional to the speed of loop extrusion *v* and independent of other parameters, indicating unobstructed loop extrusion by LEFs. In the dense regime, dynamics slows down drastically and the autocorrelation time exceeds *τ*, showing that the loop structure persists after multiple rounds of LEF exchange. In the dense regime, the autocorrelation time normalized by *τ* scales as 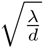

**Figure S2:**
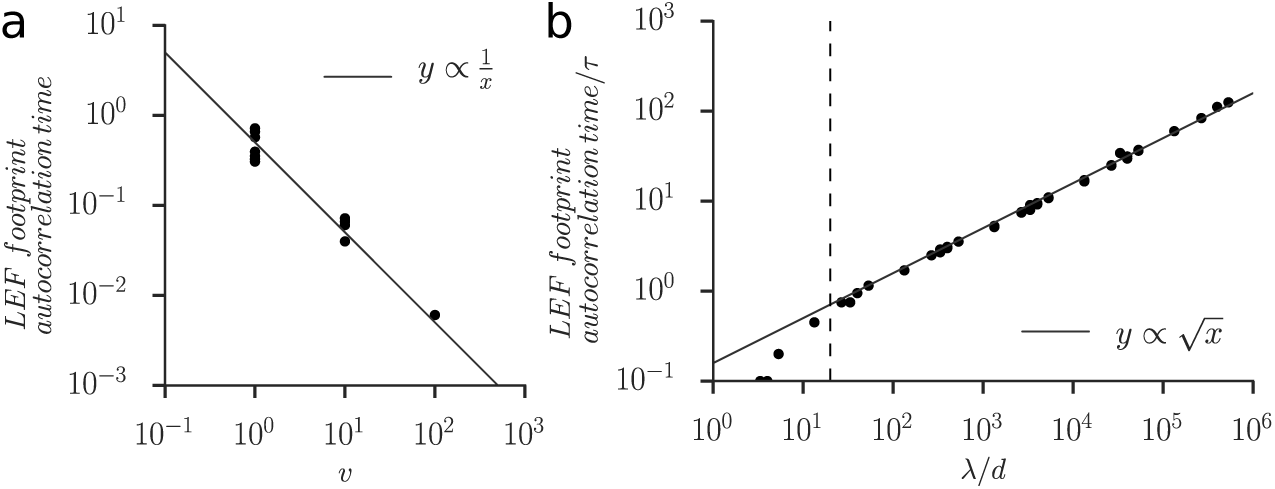
The autocorrelation time of LEF footprints in the sparse (a) and dense (b) regimes. In the sparse regime, we varied *N* between 10 and 400 LEFs, *τ* between 10 and 100, and *v* between 1 and 10. In the dense regime, *N* was varied between 100 and 1600 LEFs, *τ* between 10 and 10000, and *v* between 1 and 100. The vertical dashed line in (b) shows the approximate boundary of the dense regime, 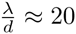.

##### 2.2 The distribution of loop lengths is exponential in the sparse regime and normal in the dense regime

The two regimes also have different statistics of loop lengths: in the sparse regime, the loop lengths are distributed exponentially; in the dense regime, the lengths are distributed approximately normally (Fig. S3).

**Figure S3:**
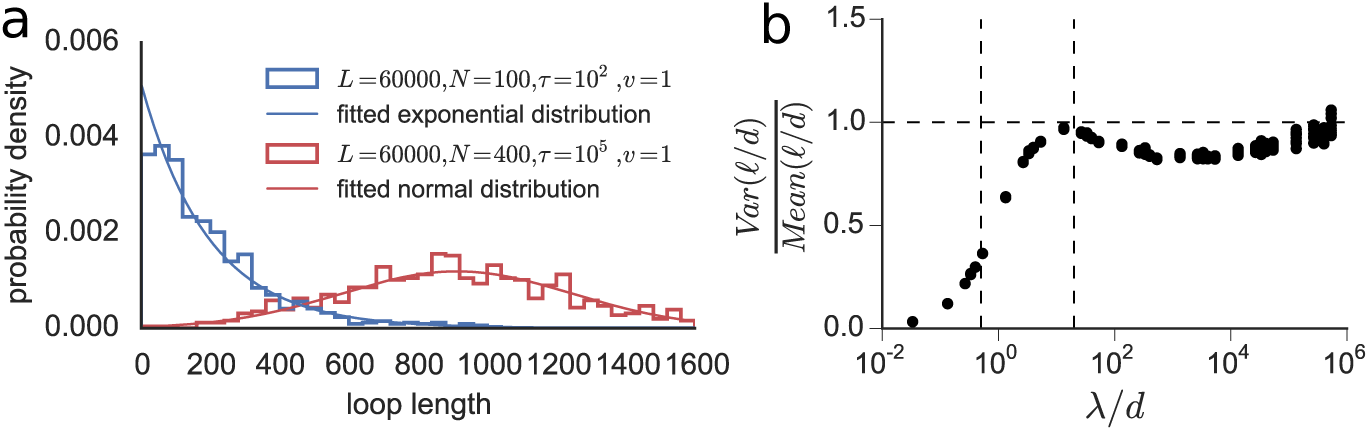
(a) Loop lengths in the sparse and dense regime follow different statistics. In the sparse regime, shown in blue, loop lengths are distributed exponentially; in the dense regime, shown in read, loop lengths are distributed approximately normally. (b) The variance-to-mean ratio of loop lengths normalized by LEF separation *d* is close to 1.0 in the dense regime.

The exponential distribution of loop lengths in the sparse regime is explained by the simple LEF dynamics. Since LEFs are separated by large gaps, they rarely block each other and extrude loops continuously throughout their residence time on the chromosome. Therefore at every moment of time, the length of a loop is proportional to the amount of time passed since its LEF bound to the chromosome. In the theory of renewal processes this amount of time is called *age* and, like the residence time of LEF, it is distributed exponentially around its mean of *τ* [4]. Thus, the lengths of loops in the sparse regime also become exponentially distributed with the average length of *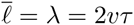*.

In the dense regime, the distribution of loop lengths can be approximated by the normal distribution, with the mean and the standard deviation that depend on the parameters of the system. Interestingly, the variance-to-mean ratio of loop lengths in units of LEF separation *d* is very close to unity in the dense regime. Our theory presented below explains why the distribution of loop lengths *ℓ* has a non-zero peak and predicts its approximate location, but it cannot predict the exact analytical form of this distribution nor its width.

##### 2.3 Gaps disappear exponentially with *λ/d*

The simple structure of loops in the sparse regime allows us to explain the observed dependence of loop coverage 1 *- g* on 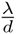 (where *g* is the portion of gaps, i.e. the portion of chromatin fiber that is not extruded into any loop) (Fig. S4). When we increase the number of LEFs *N*, while keeping *L*, *v* and *τ* fixed, the portion *g* of gaps should decrease. This dependence is, however, not linear: as more chromatin fiber is extruded into loops, it becomes increasingly likely for LEFs to form nested loops (i.e. bind within already extruded loops) and thus not contribute to the overall loop coverage. Therefore, every new LEF in the system that lands in a gap between loops increases loop coverage by 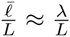 (i.e. reduces the portion of gaps *g* by the same amount). The chance of a LEF landing in a gap is *g*, which gives us a simple differential equation:

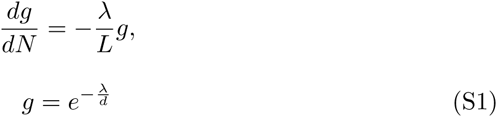

Comparison with the simulations (Figure S4) shows that this solution captures the transition between the sparse and the dense regimes, but noticeably underestimates the portion of gaps as we approach the dense regime, 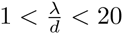 This discrepancy is due to the growing difference between *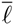* and *λ* in the dense regime (see below).

**Figure S4:**
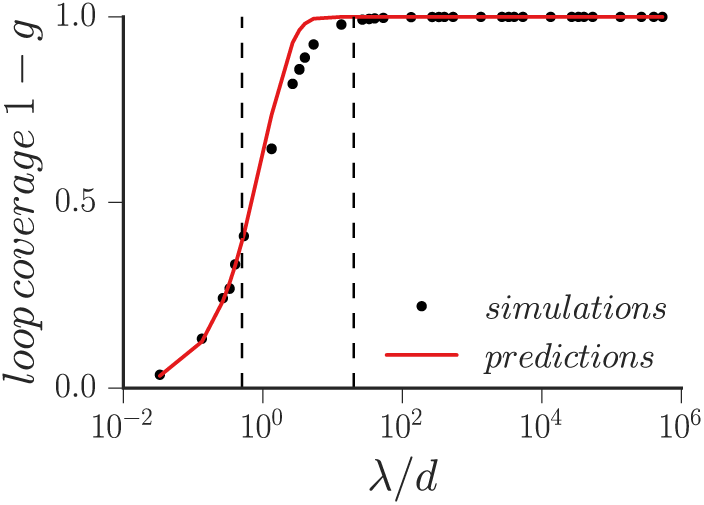
The comparison of the theoretically predicted loop coverage with the results of the simulations. The vertical dashed lines show the approximate boundaries of the sparse and dense regimes.

#### 3 The theory of self-organization of loop arrays in the dense regime

##### 3.1 In the dense regime, loops have stable lengths and fluctuating numbers of reinforcing LEFs

At *λ ≫ d*, gaps between loops disappear and the system transitions into the *dense* regime. The condition *λ ≫ d* means that LEFs can potentially form loops that are much larger than the amount of chromatin available to each of them, so the size of the extruded loop become limited by collisions between LEFs. Also, because there are no gaps between loops, LEFs that rebind to the chromatin always start new loops within other already existing loops. Finally, as we show in the main text, in the extreme dense regime loop branching becomes increasingly rare and the majority of LEFs just stack on top of each other, forming reinforced loops. Thus, LEFs in the dense regime fold chromatin into an array of consecutive reinforced loops. The length *ℓ* of each reinforced loop is relatively stable: supported by multiple LEFs, it does not disappear or shrink, when some of them unbind, but it cannot grow either, because its LEFs are blocked by the neighbors. Conversely, the number of LEFs *n* in each reinforced loop constantly fluctuates: the loops constantly lose LEFs due to their unbinding and receive new LEFs that rebind to the chromosome from the solution. Below we describe the fluctuations of loop structure in the dense regime and show how they lead to a globally stable steady state.

##### 3.2 The number of LEFs in a loop is approximately Poisson-distributed around *ℓ/d*

Let us derive the distribution of the number of LEFs *n* supporting a single reinforced loop of length *ℓ*. The loop loses LEFs due to their dissociation from the chromatin fiber. Since individual LEFs unbind with a rate 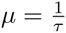, the overall rate of LEF loss depends on *n* and equals

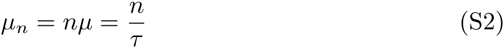

The loop also receives a flux of incoming LEFs that bind back to the chromosome from the solution. In our simple model, LEFs rebind to random sites and thus have a chance 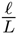 of landing within the chosen loop. As a result, the body of the loop serves as an antenna: the larger the loop is, the more LEFs it receives. At every moment of time, LEFs unbind and immediately rebind to the chromosome with the rate *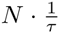*, so that the rate *r* of LEF binding to the chosen loop equals

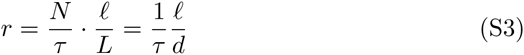

This stochastic gain and loss of LEFs produces fluctuations of the number of LEFs in the loop. The equations S2 and S3 allow us to find the probability *p_n_* for the loop to have *n* LEFs. In a population of loops of equal length *ℓ*, the fraction *p_n_* of loops supported by *n* LEFs changes over time because of three factors: a) these loops gain or lose LEFs with a combined rate *r* + *μ_n_*, b) loops with *n* − 1 LEFs gain LEFs at rate *r* and c) loops with *n* + 1 LEF lose LEFs at rate *μ_n_*+1. The dynamics of the fraction *p_n_* is then described with:

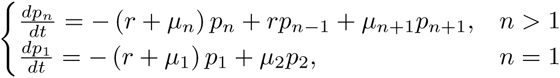

The loss of the last LEF is irreversible and loops without LEFs disappear. In order to find a quasi steady state distribution of *p_n_* described by 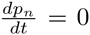, we have to ignore this fact for now and set *μ*_1_ = 0. This gives us the following system of equations [5]:

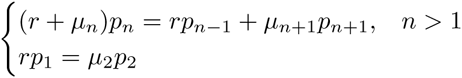

The solution of this system shows that the number of LEFs *n, n ≥* 1 in a loop is Poisson-distributed:

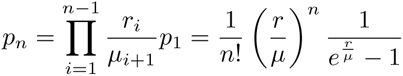

We can relate this distribution to the size of the loop and density of LEFs using the expressions (S3) and (S2):

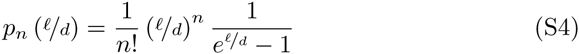

The average number of LEFs *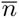* in a loop then equals:

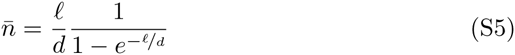

This expression highlights the importance of the length scale *d*: the length of a loop expressed in units of LEF separation *d* is approximately equal to the average number of LEFs in this loop. Below we will often use the loop length normalized by *d*:

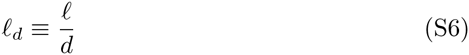

such that:

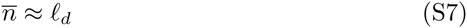

##### 3.3 The lifespan of reinforced loops increases exponentially with their length

The derived distribution (S4) of the number of LEFs per loop allows us to estimate the average lifespan of a loop *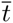*. Loops get disassembled when they lose their last LEF. Therefore, the rate of loop death can be estimated as:

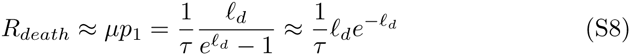

And the average lifespan *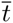* of a loop of length *ℓ_d_* is

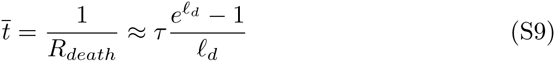

The asymptotic dependence of *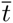* on the length of the normalized loop length *ℓ_d_* is then:

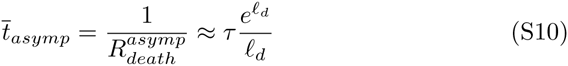

The equation (S10) reveals that the lifespan of a loop grows almost exponentially with its length. Thus, reinforcement makes longer loops essentially immortal: for example, loops with length *ℓ_d_* = 6.5 live 100 times longer than individual LEFs, and those with *ℓ_d_* = 9 live 1000 times longer. However, the effects of reinforcement are significant only for longer loops (*ℓ*_*d*_ *≳* 3.5) with the lifespan extension to 10*τ* and more.

The simple functional form of Eq. (S10) allows us to use it in analytical calculations. However, in its derivation we assumed that loop death does not perturb the distribution of *n*, which is not necessarily true. In chapter (4.1), we will derive the expression for *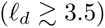* without this assumption and show that our estimates (S9) and (S10) are, in fact, very accurate.

##### 3.4 Loops in the dense regime stochastically divide in two

Loops in the dense regime divide when two LEFs land within the same loop and extrude two separate loops instead of stacking on top of one another. Both daughter loops then start receiving reinforcing LEFs, thus, cutting the supplies off the mother loop. This causes the mother loop to disassemble on the timescale of several *τ* (for more accurate estimate, see Chapter (4.4)) and the two daughter loops take its place. This process creates new loops on the chromatin in the dense regime.

We can estimate how the rate of loop division depends on the parameters of the system. LEFs land into a loop of size *ℓ* with a rate of 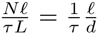. A loop divides when the second LEF lands before the first LEF has fully expanded to the loop borders, within a time window of *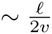*. Thus, the rate of loop division scales as:

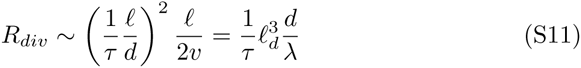

However, these two LEFs can still stack on one another. In fact, as the first LEF extrudes a bigger and bigger loop, there is less chance for the second LEF to land outside of its loop. Thus, a more accurate general expression for *R*_*div*_ should look like:

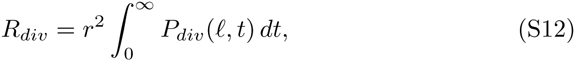

where *P*_*div*_(*l, t*) is the probability for two LEFs to divide a loop of length *ℓ* if they land with a time delay *t*.

The loop splits if the second LEF lands outside of the loop extruded by the first. Integrating over all possible positions of the two LEFs, *x*_1_ and *x*_2_, we get:

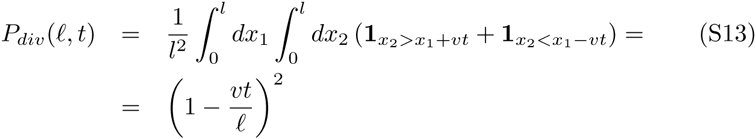

Here, **1**_*condition*_ is the indicator function, which equals 1 is the condition is true and 0 otherwise. The expression for *R*_*div*_ is then:

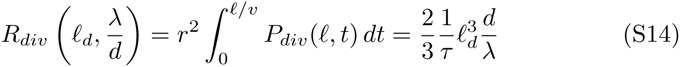

In other words, stacking of LEFs decreases the rate of loop division by 2/3.

##### 3.5 The balance between loop death and division gives rise to the steady state of the dense regime

In the steady state of the dense regime, the average length *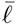* of all loops in the system stays approximately constant over time. This implies that the number of loops on the chromosome is also constant, therefore, the global rate of loop creation through division should be equal the global rate of loop death. Assuming that all loops in the system have the same length *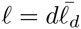*, we get the steady state condition:

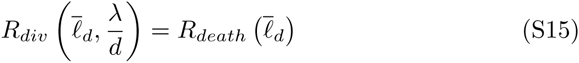

Plugging in the previously obtained estimates for *R*_*death*_ (S8) and *R*_*div*_(S14), we get:

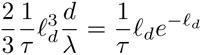

The solution of this equation defines us the average length and the number of LEFs per loop in the steady state:

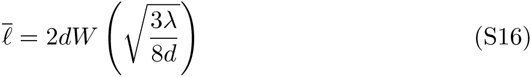

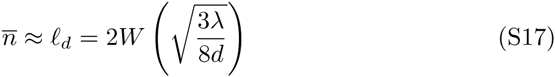

Here, *W* (*x*) is the Lambert W function, defined as the solution of *W* (*x*)*e*^*W*^ ^(*x*)^ = *x* For 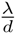 varying between 20 and 10^6^, we can use the approximation *W* (*x*) *≈* 0.3 + ln(*x*) *-* ln(ln(*x*)):

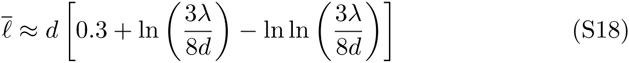

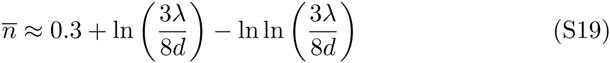

Our theory also explains why the steady state is globally stable. The average steady state loop length *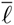* (S16) is located at the intersection of the curves *R*_*death*_(*l, d*) (S10) and division *R*_*div*_(*l, λ, d*) (S14) (Fig. S5); loops longer than *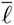* are more likely to divide into smaller loops, loops shorter than *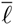* are more likely to die and let their neighbors grow. Thus, the scalings of *R*_*death*_(*l, d*) (S10) and division *R*_*div*_(*l, λ, d*) (S14) focus the distribution of the loop sizes around the average value *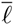*, which explains the approximately normal distribution of loop sizes shown on Fig.(S3).

**Figure S5:**
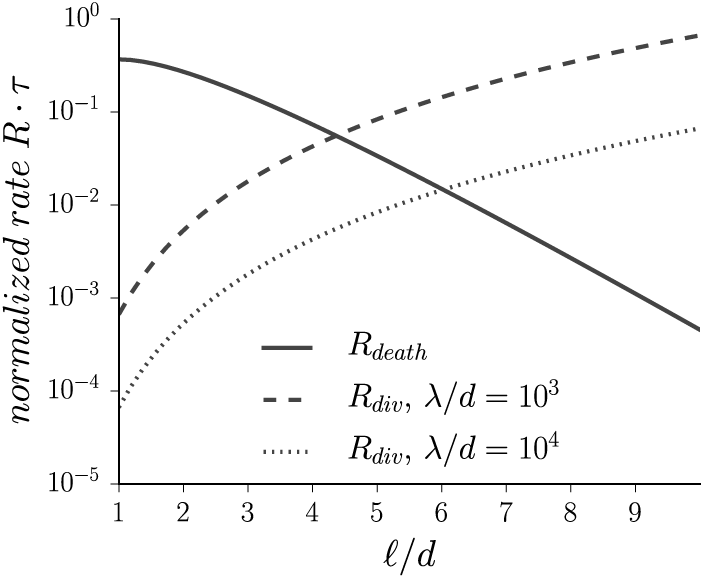
The scalings of *R*_*death*_ and *R*_*div*_ with the loop length *ℓ*. The average loop length at the steady state is located at the intersection of the two curves. The difference in the derivative signs of *R*_*death*_ and *R*_*div*_ provides global stability of the steady state.

The equation (S16) is approximate and thus can be not accurate enough. For the applications requiring ∼1% precision, we used a 7-th degree polynomial expression which was fit to 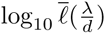 observed in the simulations with 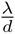 in the range (10*-*^1.5^; 10^5.5^):

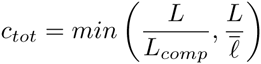

##### 3.6 Maximal lengthwise compaction is achieved on the lower border of the dense regime

LEFs fold a chromosome into a system of consecutive loops and thus dramatically reduce its length (Fig. S6). A biologically important question is what amount of LEFs would maximize lengthwise compaction of the chromosome, given their microscopic properties.

The length of a chromosome compacted by LEFs, *L*_*comp*_, has two components: (a) the combined widths of loop bases and (b) the length of gaps between loops (Fig. S6). A loop base has a width of *at least* the thickness of chromatin fiber, *a* (in our model, it equals 1 site or ∼10 nm), so that the minimal estimate for *L*_*comp*_ is:

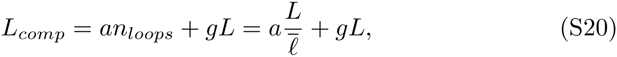

The coefficient of lengthwise compaction is then given by

**Figure S6:**
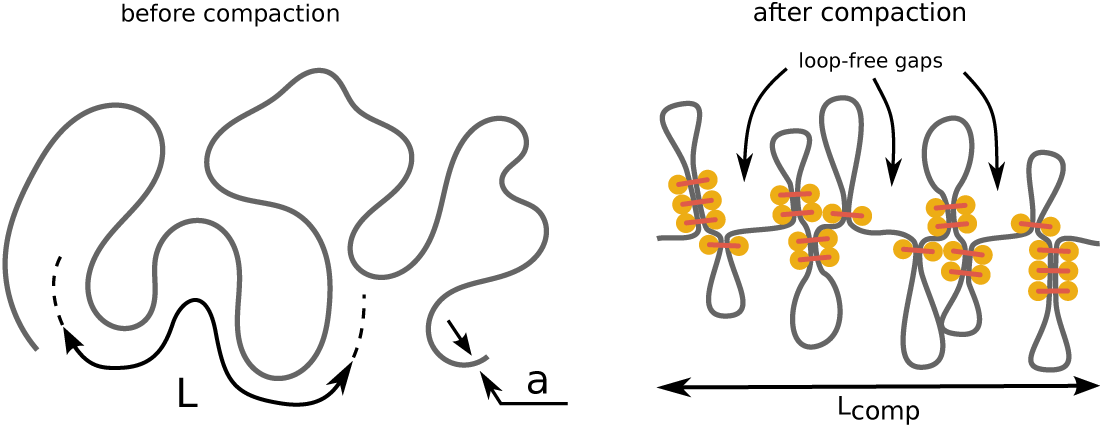
Figure S6: LEFs drastically reduce the length of a chromosome by folding it into a system of consecutive loops. The length of a compacted chromosome is determined by the combined length of the loop bases and the length of gaps between loops.

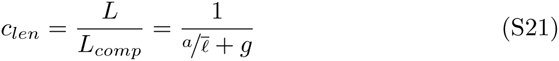

The portion of loop-free gaps *g* decreases almost exponentially with the number of LEFs as the system approaches the dense regime (see Section 2.3). On contrary, the combined length of loop bases is proportional to the number of loops in the system and grows almost linearly with the number of LEFs. Therefore, maximal compaction is achieved at the lower boundary of the dense regime, where the gaps disappear, but the number of loops is still low. Simulations confirm our reasoning (Fig.S7), but show that the exact location of the optimum depends slightly on *λ*. This dependence is caused by the presence of exponentially small residual gaps that prevent the system from achieving extremely high degrees of compaction.

##### 3.7 The maximal degree of total compaction depends on the length of the chromosome and the size of LEFs

The extreme values of the coefficient of lengthwise compaction can be misleading: high lengthwise compaction is achieved when the chromosome is folded into a few very large loops, so that the width of the compacted structure can be larger than its length. Formally, the maximal lengthwise compaction is achieved when the whole chromosome is folded into one loop! A more meaningful measure of compaction is the ratio of the chromosome length *L* to the maximum of its width and length, or, the coefficient of total compaction *c*_*tot*_. Since polymers folded into loop arrays naturally assume bottle-brush conformations [6] with loops extending away from the backbone formed by loop bases, we set the width of a compacted chromatin fiber to be roughly the size of individual loops 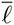:

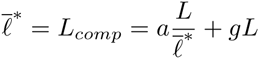

**Figure S7:**
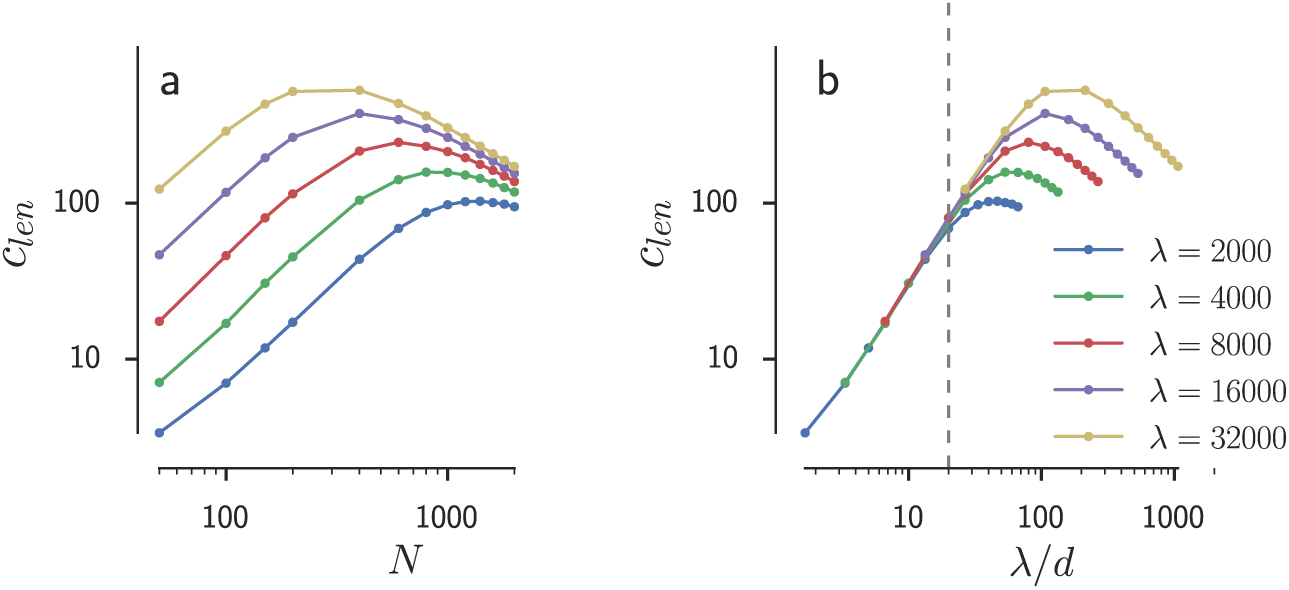
(a) Given the microscopic properties of LEFs, there is an optimal amount of LEFs that provides maximal lengthwise compaction. (b) Maximal lengthwise compaction occurs at the lower boundary of the dense regime where the gaps disappear, but the number of loops is relatively low.

Then the maximal total compaction is achieved when the width and length are equal:

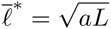

Ignoring the contribution of gaps, we can obtain the expression for the optimal loop length:

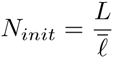

The maximal degree of total compaction is then:

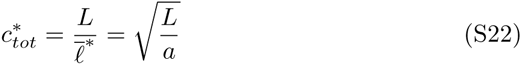

Our simulations confirm the existence of a global maximum of *c*_*tot*_ expressed by Eq. S22 (Fig. S8). The number of LEFs required to reach this maximum almost does not depend on *λ*. This is not surprising given that the average loop length in the steady state (Eq. (S16)) depends linearly on *N* and only logarithmically on *λ*. However, in order to reach the maximal total compaction, LEFs must have sufficiently large *λ*, otherwise they can not generate enough lengthwise compaction (S7).

##### 3.8 Gradual activation of LEFs speeds up convergence to the steady state

In our simulations, the steady state was achieved on long timescales, up to 10^3^*τ*. With the experimental estimates of *τ* of at least a few minutes [7], cells might not have enough time to compact their chromosomes using the mechanism described above. Below we show that gradual activation of LEFs reduces the timescales of convergence to the steady state below *τ*, thus allowing fast chromosome compaction in mitosis.

**Figure S8:**
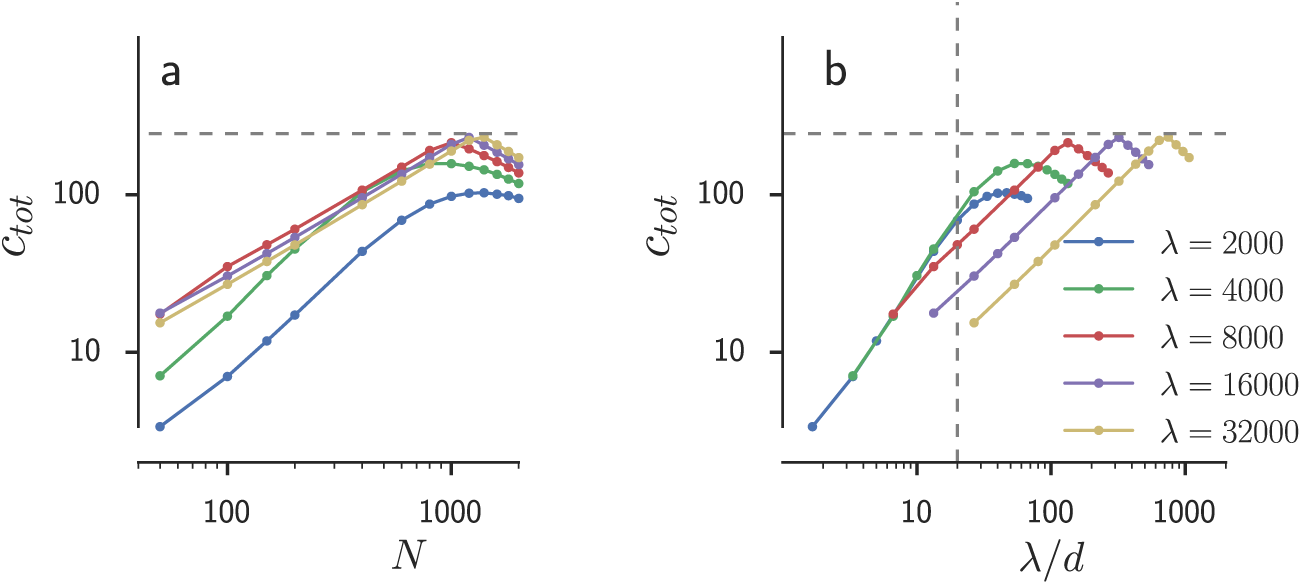
(a) The maximal total compaction and the number of LEFs required to achieve it almost does not depend on *λ*. (b) As *λ* increases, the maximal total compaction is achieved at higher values of 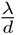. The horizontal dashed lines show the predicted maximal coefficient of total compaction 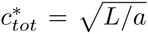 the vertical dashed line shows the lower boundary of the dense regime *λ*/*d ≈* 20.

The transition to the steady regime in our simulations is slow because we initiate the system far from the steady state. When we activate all LEFs simultaneously, they initially fold the chromosome into an array of small loops; some of these loops then slowly die and the remaining loops grow, until the average size of a loop reaches the steady state value. We can achieve a faster approach to the steady state if by initial activation of a small fraction of LEFs or if by their gradual activation.

By choosing a proper number of LEFs to activate, we can adjust the length of the initially extruded loops to be equal the steady state length. After initial loops are formed, we activate the rest of the LEFs. In this scenario the system will rapidly achieve the steady state because this second batch of LEFs will reinforce the loops formed by the first batch. The required number of initially activated LEFs is then given by the condition 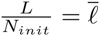

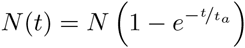

These LEFs would form an array of consecutive loops over the period of time 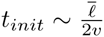, after which we could add the rest of the LEFs.

The same result can be achieved using a more realistic scenario where all LEFs are gradually (stochastically) activated with the activation rate 1/*t_a_*. The number of active LEFs in a system at time *t* then equals:

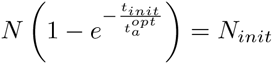

The optimal activation period *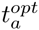* can be found by comparing this approach with the two-step activation scheme, so that by the time *t*_*init*_ the system has *N*_*init*_ active LEFs:

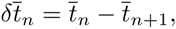

which gives us the following expression for optimal activation time *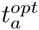*:

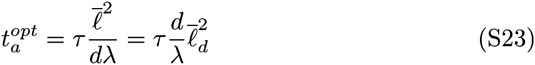

In order to confirm Eq. (S23), we simulated stochastic activation of LEFs in a wide range of parameters and found the optimal LEF activation times *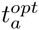*. These values of *t_a_* provided the fastest convergence to the steady state, as measured by the root mean square deviation of the loop length trajectory *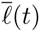* from the steady state value. The simulations showed that the expression (S23) captured the major mechanism behind the optimal LEF activation (Fig. S9).

**Figure S9:**
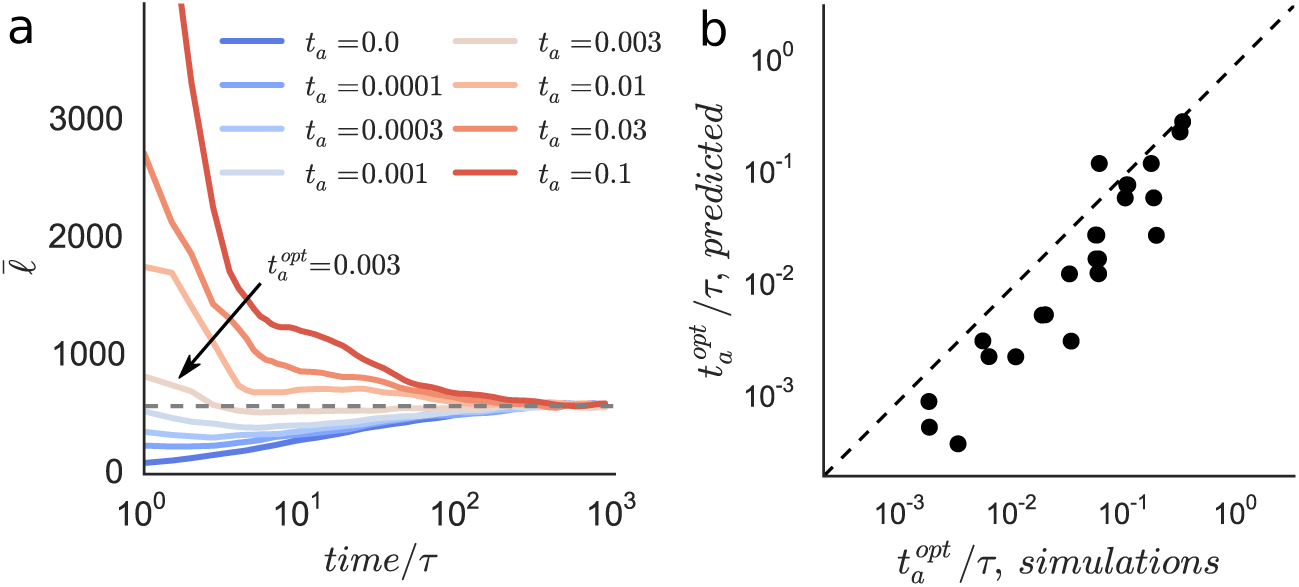
(a) Stochastic activation of LEFs with a rate 1/*t_a_* speeds up convergence of the average loop length 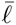 to its steady state value (the gray dashed line). The optimal activation delay *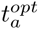* minimizes the root mean square deviation of the curve *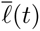* from the steady state value. (b) The values of *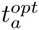* predicted with Eq. (S23) agree reasonably well with the optimal LEF activation times found in simulations across a wide range of system parameters.

The systematic discrepancy between the optimal activation times in simulations and those predicted by Eq. (S23) is due to two major factors:

a. we assumed that the LEFs activated after formation of a loop array serve only to reinforce the existing loops. However, these LEFs also divide the existing loops via the mechanism described in Chapter 3.4 and thus increase the number of created loops.
b. Some portion of the initially established *N*_*initial*_ loops dies before a loop array get established, thus decreasing the number of created loops.

Finally, it is important to note that we only considered convergence of the mean loop length 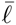; convergence of the other moments of the distribution of loop length might take a different amount of time.

### 4 Corrections to the theory of self-organization of loop arrays

#### 4.1 The theory of stochastic processes provides an exact expression for the lifespan of loops

The estimate of the rate of loop death (S8) involved several approximations and thus can potentially be inaccurate. In the next four chapters 4.1-4.4, we will use the theory of stochastic processes to derive an accurate model of loop death and estimate the precision of (S8).

We model the fluctuations of the number of LEFs *n* supporting a loop with the stochastic immigration-death process: the loop receives a steady flux of incoming LEFs (immigration), but each of them lives for a finite period of time (death). This process has been extensively studied in the literature [5] and we adapt the existing derivations to our model.

In the language of immigration-death processes, the death of loops upon loss of all LEFs corresponds to an adsorbing state at *n* = 0. Then the average lifespan of a loop is defined as the mean time to absorption into the state *n* = 0 and depends on the initial number of LEFs *n*. The lifespans *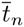* for different *n* are related to each other: a loop with *n* LEFs keeps the same *n* for an average period of 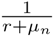 and then either receives an extra LEF with a probability *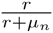* and lives for another period *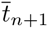* or loses a LEF with a probability 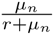 and lives for *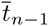* (*μ_n_* and *r* were defined in Eq.(S2) and Eq.(S3)). This allows us to relate different *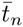* with a system of equations:

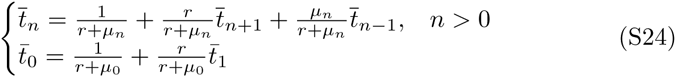

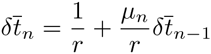

This system can be converted into a recursive equation using a new variable *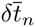*

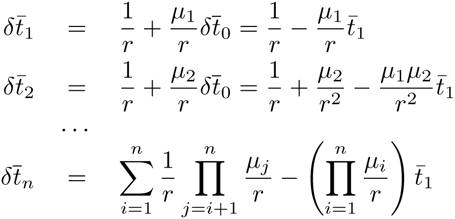

such that *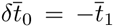* and 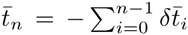 Plugging *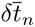* into the system of equations (S25), we obtain a recursive relation

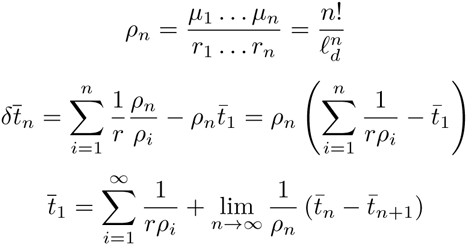

Then,

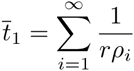

Defining an auxiliary variable *ρ_n_*, we get:

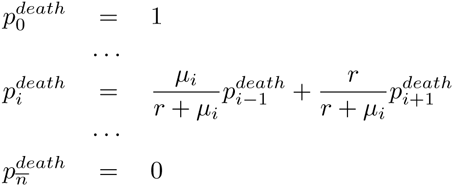

The second term is zero because 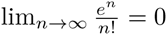, therefore

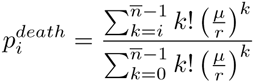

This finally allows us to find 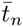:

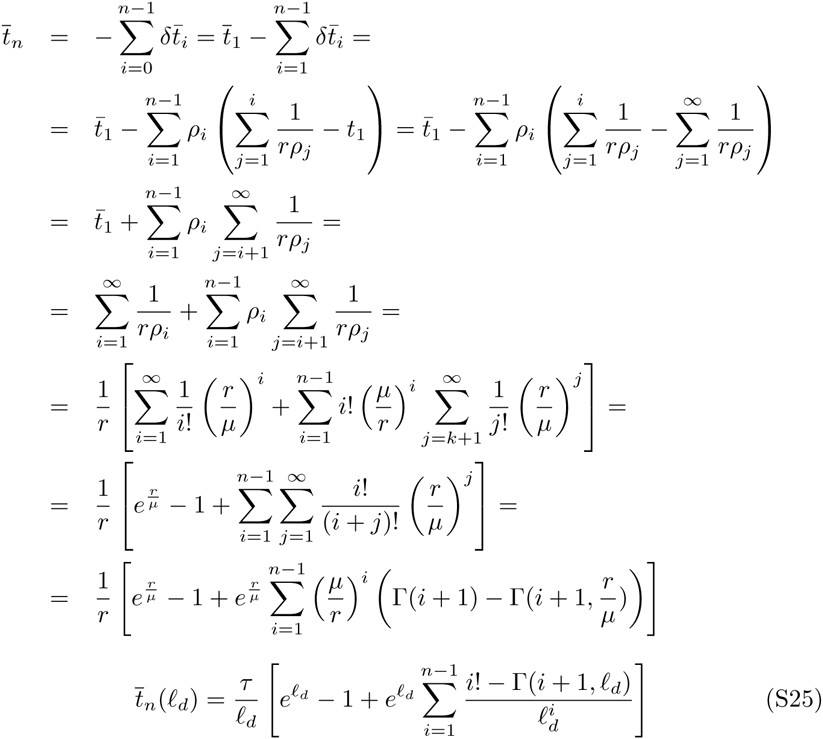

where γ(*a, x*) is the upper incomplete Gamma function. Interestingly, the previously obtained expression for *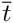* (S9), in fact, equals the average lifespan of a loop with a single LEF:

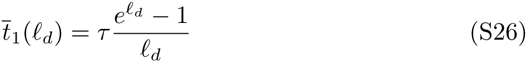

The expression (S25) is too bulky to be used in analytical derivations or provide any new intuitive understanding. Below we show that most of the terms in (S25) in fact can be dropped to obtain a compact, yet accurate approximation of *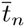*.

#### 4.2 Loops supported by a single LEF can die immediately

The Eq. (S25) gives us the average loop lifespan, but does not tell anything about the distribution of *t_n_*. In order to find it, we simulated 10^5^ stochastic immigration-death processes with *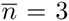* with *n*_0_ = 1 and *n*_0_ = 3 initial LEFs (Fig. (S10)). Both distributions had exponential tails, but the distribution of 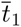 had an additional peak around zero (Figure S10). The normalized tails of the distributions for the two initial conditions perfectly matched at *t*_*n*_ *≥* 3*τ*, indicating that the starting number of LEFs did not affect the later stages of loop dynamics.

**Figure S10:**
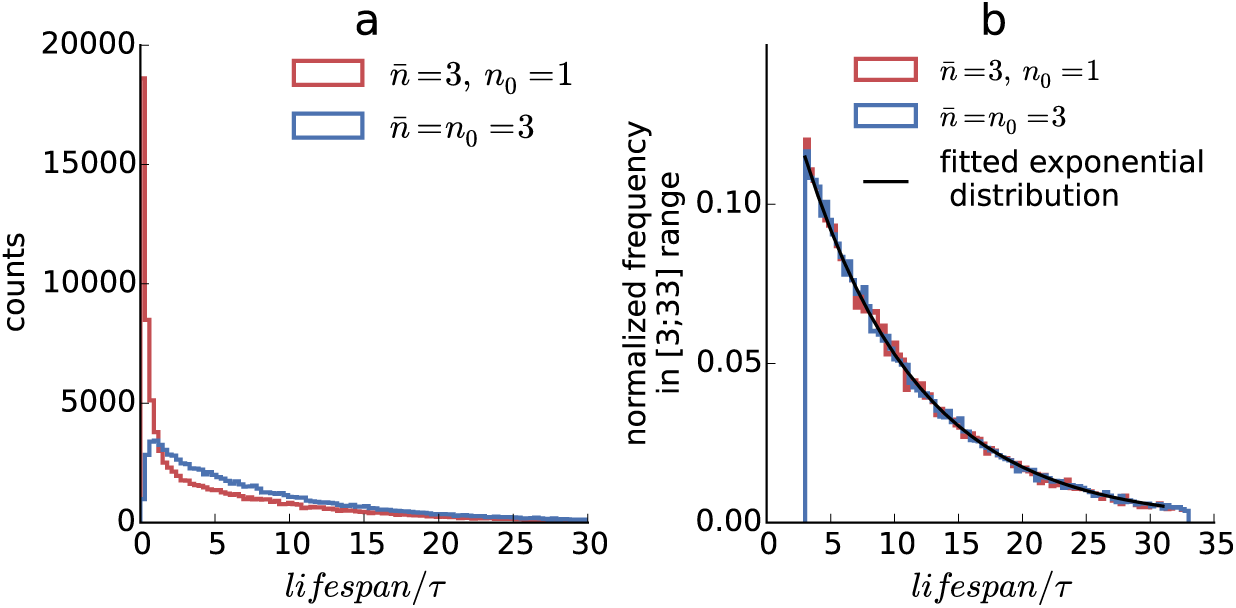
The distribution of the lifespans of immigration-death processes with different initial conditions. (a) The overall distribution; (b) the normalized lifespan frequency for the processes that survived the initial period *t*_*n*_ *<* 3*τ*.

The increased frequency of short lifespans of single-LEF loops is caused by stochastic fluctuations: with some probability these loops immediately lose their only LEF and disassemble before receiving any reinforcing LEFs. A rough estimate for the probability of such immediate death is:

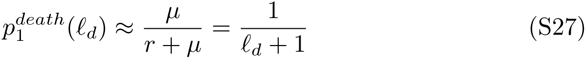

We can find a more accurate expression for *p*^*death*^ using the theory of immigration death processes. If we define immediate death more generally as an event when a loop dies before reaching its fully reinforced level *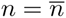*, we get an immigration-death process with two adsorbing boundaries at *n* = 0 and *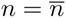*. Then the probability *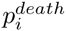* that a loop with *i* LEFs will die before accumulating *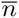* LEFs, obeys the following system of equations:

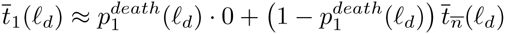

And the solution is:

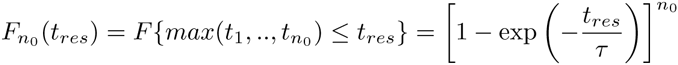

Particularly, a loop formed by a single LEF dies before accumulating *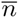* LEFs with the probability

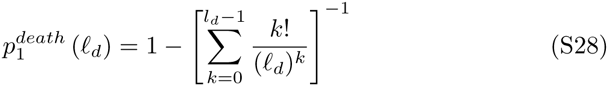

#### 4.3 Accounting for immediate death gives a simple and accurate estimate for the rate of loop death

The analysis above suggests two different processes lead to loop death: (a) immediate death before full reinforcement, which probability depends on the initial number of LEFs *n*_0_, (b) exponential death of fully reinforced loops, which rate is independent of *n*_0_. This allows us to find a compact estimate for 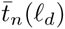 Roughly, a single-LEF loop either dies immediately with a chance *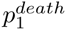* or quickly accumulates *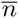* LEFs and lives for *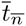*:

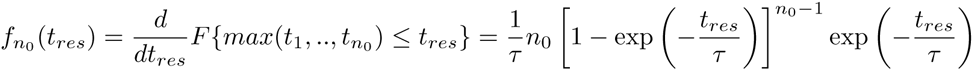

Then the lifespan of a loop with *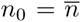* LEFs is given by:

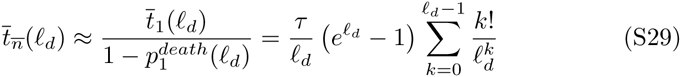

This formula can be generalized for an arbitrary initial number of LEFs *n*_0_, 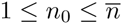

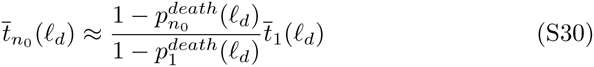

Now we can compare the accuracy of the expressions (S25), (S10) and (S29). We simulated the dynamics of LEF number *n* using a stochastic immigration-death process with different average numbers of LEFs *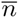*, *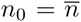*. Simulations at each value of *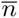* were repeated 1000 times to obtain enough statistics (Figure (S11)).

**Figure S11:**
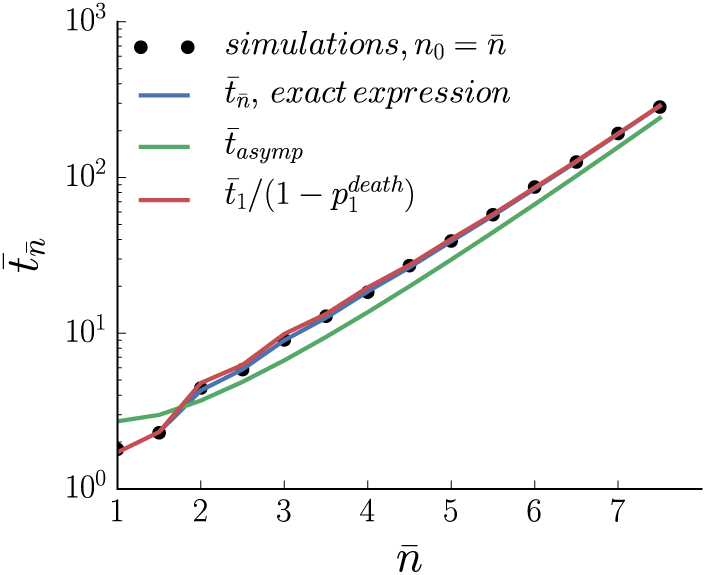
The average lifespan of a loop can be predicted accurately with analytical expressions. The black dots show the results of simulations, the predictions with analytical expressions (S25), (S10) and (S29) are shown in blue, green and red, correspondingly.

The simulations show that the asymptotic expression (S10) significantly overestimates the lifespan 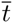 of short loops (*ℓ*_*d*_ *<* 2) and underestimates 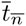 for longer loops. On contrary, the approximate expression (S29) is as accurate as the exact expression (S25) and thus can be used instead.

#### 4.4 The lifespan of a loop after interruption of reinforcement is approximately Gumbel-distributed

In our theory of loop division, we assumed that “mother” loops die soon after interruption of reinforcement. In this chapter we support this assumption with analytical derivations and obtain the full distribution of residual lifetimes of loops.

A loop exists as long as it has at least one LEF at its base. Therefore, its residual lifespan *t*_*res*_ after interruption of reinforcements is given by the maximum of the residence times of its *n*_0_ LEFs. This allows us to write the cumulative distribution function (CDF) of *t*_*res*_ as the CDF of the maximum of *n*_0_ exponentially distributed lifetimes of individual LEFs:

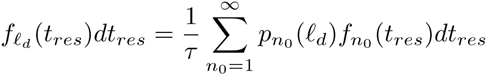

The probability density function of *t*_*res*_ is then given by:

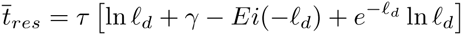

We can rephrase this expression in terms of the known loop length *ℓ_d_* by summing over all possible LEF number *n*_0_:

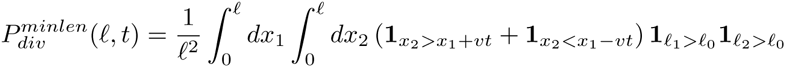

Since the number of LEFs *n*_0_ is roughly Poisson-distributed around *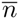* (Eq. (S4)), we get

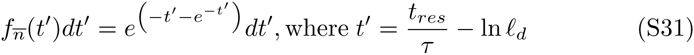

The average lifespan of a loop 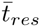 after interruption of LEFs is then

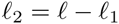

Here, *γ* ≈ 0.577 *…* is the Euler–Mascheroni constant and *Ei*(*x*) is the exponential integral. At larger values of *ℓ_d_* this expression can be further simplified to:

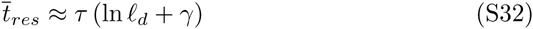

The expression (S31) looks exactly like the Gumbel distribution, with the only difference that it is defined on *t’* ∈ [– ln *ℓ_d_*; ∞), while the Gumbel distribution is defined on the whole real line. The fact that the lifespan of a stack of LEFs is roughly Gumbel-distributed is not surprising: the residual lifespan of a loop equals the maximum of *n*_0_ exponentially-distributed lifespans of individual LEFs and the Gumbel distribution is the limiting distribution of a maximum of *n* exponential numbers when *n → ∞*.

#### 4.5 Selection for the minimal daughter loop size slows down loop division

In the next three chapters 4.5-4.7, we will analyze the inaccuracies of our simple model of loop division and will come up with better estimates for *R*_*div*_.

The first major correction to Eq. (S14) is due to the fact that both daughter loops must be large enough to survive and receive stable reinforcements. A more accurate estimate of *R*_*div*_ therefore must discard the configurations of daughter loops where one of them is too small to be viable. The condition on the minimal size of the loop then can be plugged into the expression for the probability of division *P*_*div*_ (S13):

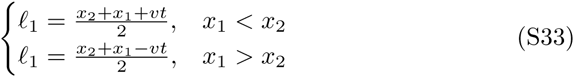

Here, *ℓ*_0_ is the minimal length of a daughter loop, *ℓ*_1_ and *ℓ*_2_ are the lengths of the fully-extruded daughter loops, defined as:

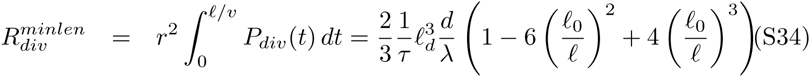

This gives us the following expression for the rate of loop division:

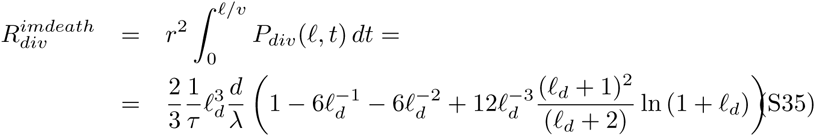

We choose the minimal loop size to be *ℓ*_0_ = 3*d*, since it is the minimal loop size required for an order of magnitude increase of the lifespan.

#### 4.6 Immediate death of daughter loops slows down loop division

Eq. (S14) also ignores the fact the newborn daughter loops have only one LEF and thus can die before becoming fully reinforced. This can happen even to large loops with the length *l > l*_0_ and thus this effect is different from the one considered in the previous chapter.

The expression for the probability of successful division (S13) can be modified to take into account a chance of immediate death:

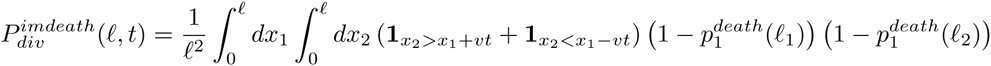

The rate of loop division then can be calculated analytically if we truncate the expression for *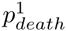* at the second term:

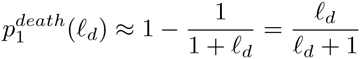

#### 4.7 Loop size selection and immediate death affects daughter loops independently

The most accurate expression for *R*_*div*_ should account both for the size selection and immediate death of daughter loops:

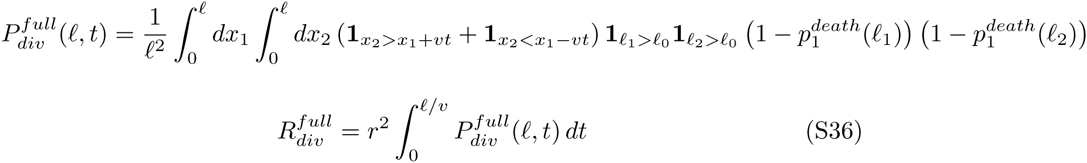

This expression does not have a short analytical form and has to be calculated numerically.

We compare the accuracy of the four different expressions for *R*_*div*_ (S14), (S34), (S35) and (S36) with the results of our simulations. For every tested combination of parameters (*L, N, v, τ*), we measure the rate of loop division *R*_*div*_ in the simulations and then estimate it with the equations (S14), (S34), (S35) and (S36) using the observed distribution of loop lengths (Fig. S12). We found that both corrections for loop size selection and immediate death of daughter loops significantly improve the accuracy of predicted *R*_*div*_, with the combined expression (S36) having the highest accuracy.

**Figure S12:**
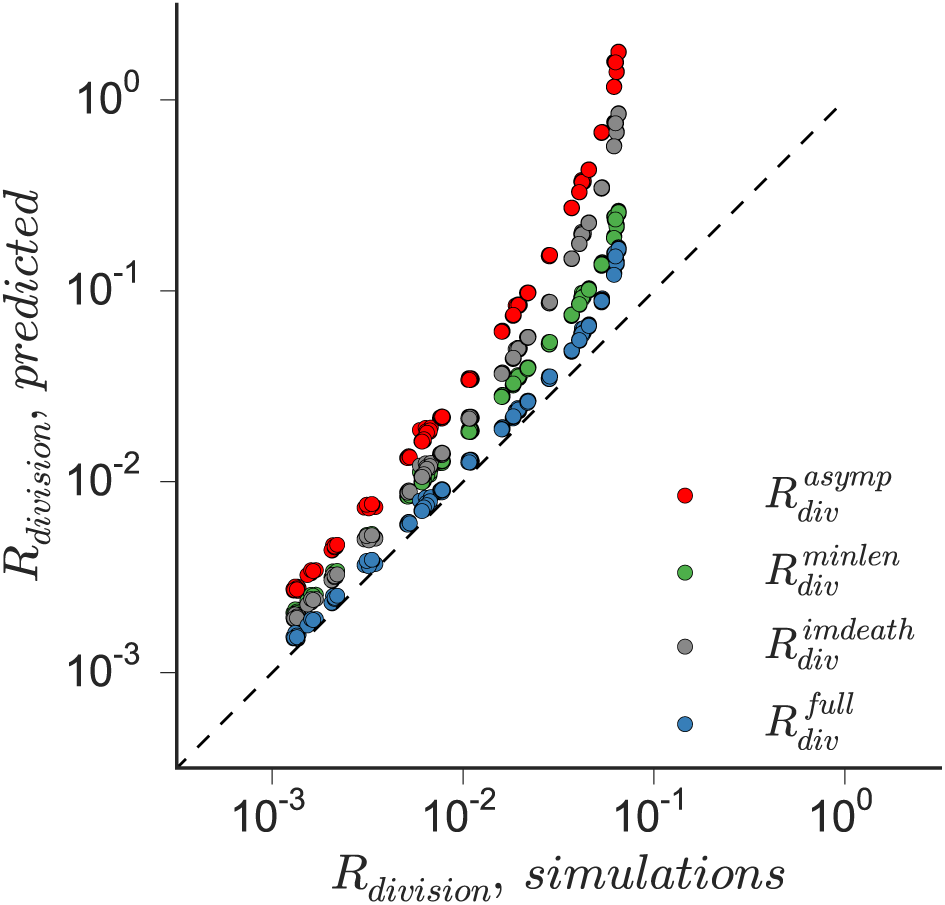
The rate of loop division in simulations can be predicted accurately using analytical expressions. The predictions with analytical expressions (S14), (S34), (S35) and (S36) are shown in red, green, gray and blue, correspondingly.

### 5 Glossary and mathematical notation

- Loop extruding factor (LEF) - a molecular machine which bridges two adjacent sites on a chromosome and then moves the binding sites in the opposite directions along the chromosome, thus extruding a loop.
- Nested loop - a loop extruded inside some other loop (Fig.S13).
- Reinforced loop - a chromatin loop supported by several LEFs closely stacked on top of one another. Alternatively, a series of nested loops, formed by closely stacked LEFs.
- Branched loop - a loop containing two or more nested loops that are not nested into one another.
- Root loop - a loop that is not nested into any other loop; includes all nested loops, if it has any.
- *L* - the length of the chromosome.
- *N* - the number of LEFs in the system.
- *τ* - the average time that a LEF stays continuously bound to the chromosome.
- *v* - the average speed with which a LEF motor translocates chromatin fiber.
- *λ* - LEF processivity, the average length of a chromatin loop that a single unobstructed LEF can extrude over its residence time.
- *d* - LEF separation, the average spacing between LEFs if they were randomly dispersed along the chromosome.
- *a* - the thickness of the fiber.
- *n* - the number of LEFs supporting a reinforced loop.
- *ℓ* - the length of a loop.
- *ℓ_d_* - the length of a loop normalized by the LEF separation.
- *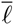* - the average length of a loop in the steady state.
- *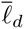* - the average length of a loop in the steady state normalized by the LEF separation.
- *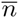* - the average number of LEFs supporting a reinforced loop in the steady state.
- *r* - the rate of LEF binding per loop.
- *μ* - the rate of LEF unbinding.
- *f* - the fraction of the chromosome contained in the gaps between the loops.
- *c*_*len*_ - the coefficient of the lengthwise compaction.
- *R*_*death*_ - the rate of death of reinforced loops.
- *R*_*div*_ - the rate of division of reinforced loops.
- *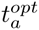* - the optimal rate of stochastic LEF activation that provides the fastest convergence to the steady state.

**Figure S13:**
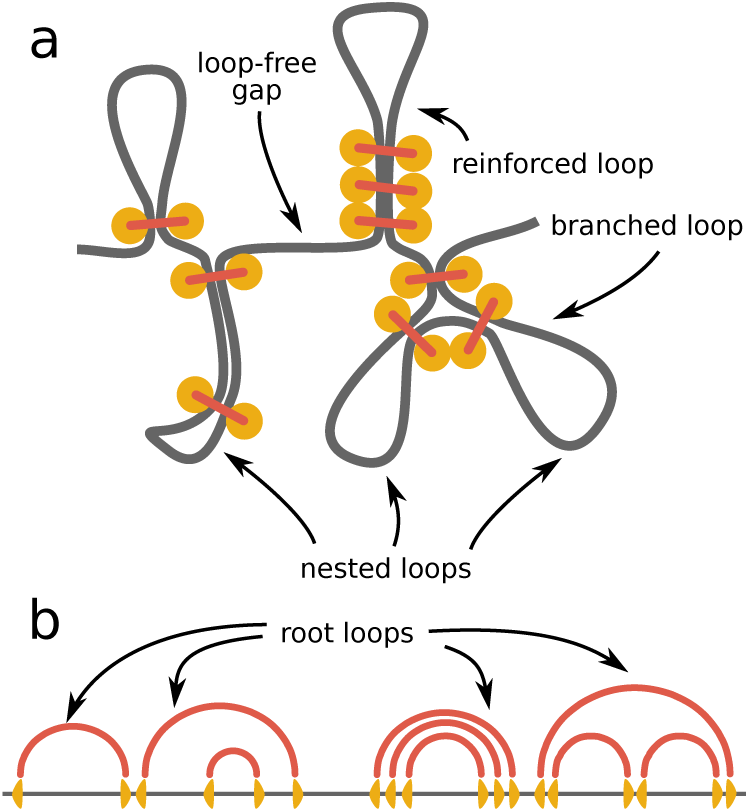
(a) The illustration of the possible loop structures formed by loop-extruding factors (LEFs). (b) The corresponding diagram of the intramolecular links formed by LEFs.

